# Mechanism Underlying Carvacrol-induced Alveolar Bone Repair via the Phosphatidylinositol 3-kinase Pathway in Periodontal Disease

**DOI:** 10.1101/2025.02.07.637017

**Authors:** Teng Nian, Li Zhaoxun, Gao Tiantian, Xiang Yanrui, Zhou Lulu, Gao Xiang

## Abstract

**Background:** Conventional antibiotic treatments do not restore the structure of periodontal tissues and can cause antibiotic resistance. This study describes the mechanism by which carvacrol regulates alveolar bone osteoblasts in periodontal disease.

**Methods:** A rat model of periodontal disease (P group) was created by ligating the first molar and injecting lipopolysaccharide (LPS). Additionally, a control model (C group) was created. The treatment models received low (L group), medium (M group), and high (H group) doses of carvacrol hydrogel. In vitro, rat osteoblast cells were divided into the C, P, L, M, H, and carvacrol + LY294002 (Car+LY group) groups. Immunohistochemical staining analyzed collagen type I and runt-related transcription factor 2 expressions. Western blot analysis detected the phosphatidylinositol 3-kinase (PI3K)/protein kinase B (AKT)/glycogen synthase kinase 3 beta (GSK-3β) pathway-related and osteoblastic proteins. Quantitative reverse transcription polymerase chain reaction measured the expression of inflammatory factors and osteoblastic proteins. The alkaline phosphatase (ALP) colorimetric kit and Alizarin Red S staining kit assessed the osteogenic ability. Detect the expression of COL1 in osteoblasts by immunofluorescence (IF). Transmission electron microscopy detected cell apoptosis.

**Results:** Carvacrol hydrogel alleviated periodontal symptoms, upregulated PI3K/AKT/GSK-3β pathway-related and osteoblastic proteins, and increased ALP expression and calcified nodules. However, it decreased cell apoptosis and inflammatory factors. LY294002 inhibited the PI3K/AKT pathway and decreased osteoblastic protein expression, ALP coloration, and calcified nodules.

**Conclusion:** Carvacrol hydrogel promotes the proliferation and differentiation of alveolar bone osteoblasts by activating the PI3K pathway and inhibiting inflammation-induced bone resorption. This study emphasizes the potential of carvacrol for treating periodontal diseases.

## 1. Introduction

Periodontal disease is a chronic inflammatory disease that adversely affects the health and quality of life of both humans and animals^[1]^. It is characterized by the progressive destruction of the tissues supporting the teeth, leading to alveolar bone resorption, tooth loss, and systemic complications^[2]^. The initiation and progression of periodontal disease have been strongly associated with bacterial proliferation in the periodontal environment, which triggers an inflammatory response in the gums and surrounding tissues. This inflammation can extend beyond the oral cavity, contributing to systemic health issues^[3]^.

Carvacrol—a monoterpene isolated from thyme (*Thymus vulgaris*) and oregano (*Origanum vulgare*)—has garnered attention for its wide availability, low cost, and high safety profile^[4]^. Carvacrol exhibits numerous beneficial properties, including anti-inflammatory, antibacterial, antioxidant, and bone-protective effects^[5]^. Previously, it has been demonstrated that carvacrol effectively inhibits *Porphyromonas gingivalis* and *Fusobacterium nucleatum*, two key pathogens in periodontal disease. Furthermore, carvacrol reduces alveolar bone resorption associated with periodontitis^[6]^. These properties position carvacrol as a promising candidate for the treatment of periodontal disease.

Despite the potential of carvacrol, its exact mechanisms of action underlying periodontal tissue repair remain elusive. Particularly, the role of carvacrol in regulating osteogenic processes, such as osteoblast proliferation and differentiation, is unclear. To this end, this study elucidates the ability of carvacrol to promote alveolar bone repair by inhibiting osteoclast activity and enhancing osteoblast function.

To better understand the molecular mechanisms underlying the effects of carvacrol, the role of the phosphatidylinositol 3-kinase (PI3K)/protein kinase B (AKT) signaling pathway and β-catenin in osteoblast function was investigated. The PI3K/AKT pathway is central to regulating cell growth, proliferation, migration, metabolism, and survival^[7]^. Activation of the PI3K/AKT and glycogen synthase kinase 3 beta (GSK-3β) pathways promotes osteoblast proliferation and differentiation^[8]^.

In this study, a rat model of periodontal disease was used to investigate the effects of carvacrol on periodontal tissue repair and the underlying molecular mechanisms. The findings provide a theoretical foundation for developing novel plant-derived drugs targeting alveolar bone osteogenesis. This study set out to investigate the potential of carvacrol as a novel therapeutic agent for periodontal disease. By elucidating the mechanisms by which carvacrol promotes alveolar bone repair and inhibits osteoclast activity, the present study will contribute to the development of more effective treatments for this debilitating condition.

## 2. Materials and Methods

The study protocol was approved by the Northeast Agricultural University Veterinary Review Committee (approval number: NEAUEC20230341).

### 2.1 Materials

The consumables required for the experiment are listed after the references.

### 2.2 Acquiring Animals and Establishing the Periodontal Disease Model

50 four-week-old male SPF Sprague-Dawley rats weighing between 180 g and 200 g were used, purchased from Liaoning Changsheng Biotechnology Co., Ltd. All rats, except those in the control group (C group), were fasted and weighed under anesthesia. They were fixed supine on the experimental table. First, using a blunt periodontal probe, artificial periodontal pockets were created around the bilateral maxillary first molars. Second, the necks of the bilateral maxillary first molars were ligated with 0.2 mm diameter orthodontic wire, ensuring the wire was completely submerged into the gingiva. Lipopolysaccharide (LPS) (20 μL) was injected into the pockets^[9]^. The ligature was placed for 4 weeks. Red and swollen gums, bleeding upon probing, and periodontal pockets indicated successful modeling^[10]^.

After the final drug intervention, the periodontal indexes were evaluated. Five experienced clinical teachers from the Surgical Laboratory of the College of Animal Medicine of Northeast Agricultural University and the Teaching Animal Hospital of Northeast Agricultural University were invited to participate in the evaluation.

### 2.3 Animal Grouping and In Vivo Drug Treatment

Forty rat models were randomly divided into four groups (10 rats per group) as follows:

- Periodontal disease group (P group): No additional treatment.
- Low-dose carvacrol hydrogel group (2.5 uL/mL carvacrol L group): Injected with low-dose carvacrol hydrogel.
- Medium-dose carvacrol hydrogel group (5 uL/mL carvacrol M group): Injected with medium-dose carvacrol hydrogel.
- High-dose carvacrol hydrogel group (10 uL/mL carvacrol H group): Injected with high-dose carvacrol hydrogel.

Hydrogel preparation: Pluronic F127, F68, and hydroxypropyl methylcellulose (HPMC) were dissolved in deionized water to prepare solutions with concentrations of 30%, 10%, and 2.5%, respectively. Subsequently, these reagents were thoroughly mixed in a volume ratio of F127: F68: HPMC = 7:1:1.5 to obtain a blank hydrogel^[11]^. Different volumes of carvacrol were added to the blank hydrogel to prepare three doses of carvacrol hydrogels. Previously, the administration method and hydrogel performance have been confirmed by us as an effective method of drug delivery. The injection method caused no significant damage to bone, and the related results have been published^[6]^.

Gingival bleeding index (GBI) scoring criteria^[12]^ (Table 1)

**Table 1.**
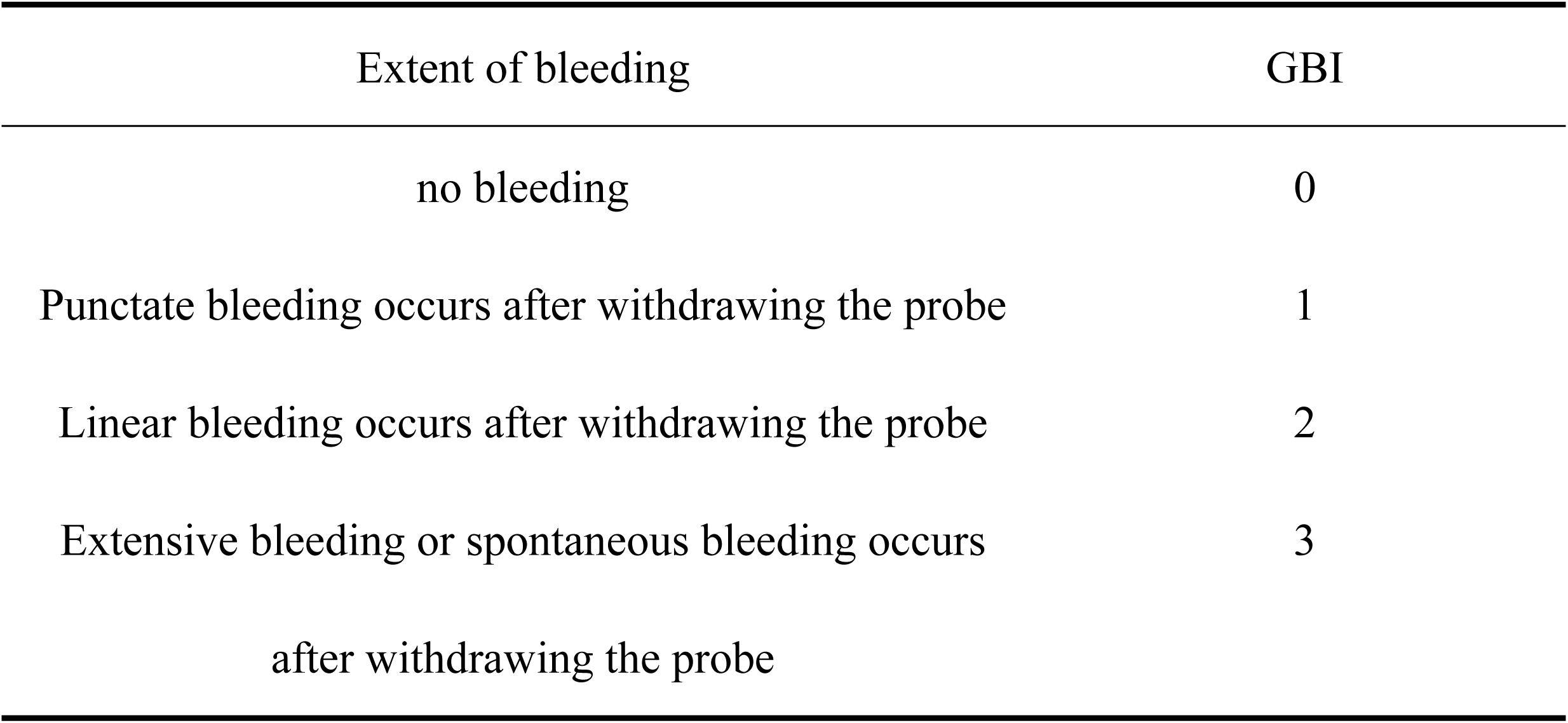
GBI scoring criteria.

Pocket depth (PD): Using a graduated periodontal probe, the distance from the gingival margin to the bottom of the periodontal pocket was measured in three areas, ensuring the probe tip remained in close contact with the tooth surface and parallel to the long axis of the tooth^[13]^.

After 4 weeks of ligature, orthodontic wires were removed. For all groups, except the C group, 50 μL of the corresponding drug was injected into the periodontal pockets twice weekly for 4 weeks. GBI and PD were measured twice weekly.

### 2.4 Micro-Computed Tomography

Four rats were selected from each group, and their maxillary alveolar bone samples were fixed and scanned using micro-computed tomography (micro-CT). Alveolar bone samples were fixed in 4% formaldehyde solution and subjected to micro-CT scanning. The scanning parameters were set as follows: tube current of 200 uA, voltage of 85 KV, scanning of the entire object, scanning resolution of 10.153741 μm, exposure time of 384 ms, and scanning angle of 180°. Samples from this experiment were submitted to an authoritative testing institution at Shanghai Jiao Tong University for analysis.

### 2.5 Histology

Maxillary tissues were isolated, fixed, decalcified, and embedded. They underwent H&E (hematoxylin and eosin) staining and immunohistochemistry (IHC). The tissues were observed under a microscope, and the images were collected and analyzed^[14]^.

### 2.6 Western Blot Analysis

Gingival tissues were ground and treated with radioimmunoprecipitation assay buffer (consisting of 1% phenylmethylsulfonyl fluoride and 1% phosphatase inhibitor). They were centrifuged at 12,000 rpm for 10 min at 4°C. The supernatant was collected, and protein concentration was measured using the bicinchoninic acid assay. Proteins were denatured with a loading buffer and phosphate-buffered saline (PBS), heated at 100°C for 10 min, separated via sodium dodecyl sulfate-polyacrylamide gel electrophoresis, and transferred to polyvinylidene fluoride membranes. After blocking with 5% non-fat milk for 2 h, the membranes were incubated overnight at 4°C with primary antibodies, followed by incubation with secondary antibodies for 2 h at 25°C. Protein bands were visualized using electrochemiluminescence reagents and imaged using a chemiluminescence gel imaging system. Western blot indicators included phosphorylated phosphatidylinositol 3-kinase (p-PI3K), p-AKT, p-GSK-3β, β-catenin and osteogenesis-related proteins, such as osteopontin (OPN), osteoprotegerin (OPG), collagen type I (COL1) and Runt-related transcription factor 2 (Runx2). The results were normalized to β-actin.

### 2.7 Quantitative reverse transcription polymerase chain reaction analysis

Total RNA was extracted using TRIzol reagent, quantified using a spectrophotometer, and reverse-transcribed into cDNA. The relative expressions of collagen type I alpha 1 chain (COL1A1), OPN, Runx2, OPG, ALP, interleukin (IL)-6, IL-1β, and tumor necrosis factor-alpha (TNF-α) in rat gingival tissues were measured using quantitative reverse transcription polymerase chain reaction (qRT-PCR), with β-actin as an internal reference. Each sample was replicated six times, and the 2^−ΔΔCt^ method was used to calculate the relative expressions.

### 2.8 Cell Culture Grouping and In Vitro Experiments

Rat osteoblast ROS17/2.8 cells were cultured in high-glucose Dulbecco’s Modified Eagle Medium supplemented with 10% fetal bovine serum and 1% penicillin-streptomycin. The experimental groups are mentioned below.

Cells were seeded at a density of 5 × 10^3^ cells/mL in 96-well plates (100 μL/well) and incubated at 37°C with 5% CO_2_. After 24 h, fresh media consisting of different carvacrol concentrations (0, 50, 100, 150, 200, 250, and 300 μM) replaced the original media. After 24 h, the original media was removed, and fresh media mixed with CCK-8 reagent (1:10 dilution) was added (110 μL/well). The plates were incubated for 2 h at 37°C with 5% CO_2_, and absorbance was measured at 450 nm using an enzyme-linked immunosorbent assay reader. A similar method was used to determine the effect of LY294002.

First part: Screening the optimal concentration for administration.

C group: No LPS or carvacrol;

P group: 1 μg/mL LPS;

L group: 1 μg/mL LPS + 50 μM carvacrol;

M group: 1 μg/mL LPS + 100 μM carvacrol; and

H group: 1 μg/mL LPS + 150 μM carvacrol.

Second part: PI3K inhibitor was added to confirm the underlying molecular mechanisms.

C group: No LPS or carvacrol;

P group: 1 μg/mL LPS;

Car group: 1 μg/mL LPS + 150 μM carvacrol; and

Car+LY group: 1 μg/mL LPS + 150 μM carvacrol + 10 μM LY294002.

### 2.9 ALP and Alizarin Red S Staining

Cells were seeded at a density of 1 × 10^5^ cells/well in six-well plates. The cells reached 80% confluence, following which the culture medium was replaced with an osteogenic induction medium. After 7 days, ALP staining was conducted^[15]^. For Alizarin Red S staining, the cells were cultured for 28 days and stained according to the manufacturer’s instructions.

### 2.10 qRT-PCR and Western Blot Analysis

The methods and indicators for qRT-PCR and western blot analysis were similar to those used for the animal experiments.

### 2.11 Electron Microscopy

First, logarithmic phase cells were seeded in culture flasks, incubated for 24 h, and divided into C, P, Car, and Car+LY groups. Second, the cells were digested with trypsin, collected, and washed three times with PBS. Third, they were fixed with 2% glutaraldehyde, post-fixed with 1% osmium tetroxide, and embedded in Spurr resin. Finally, the cells were sectioned and observed under transmission electron microscopy (TEM).

### 2.12 Immunofluorescence (IF)

Cells were seeded at a density of 1 × 10^3^ cells/well in 24-well plates and incubated for 12 h. They were divided into the C, P, Car, and Car+LY groups. These cells were fixed, blocked for 2 h at room temperature, and incubated overnight at 4°C with a COL1 antibody (1:200 dilution). They were washed twice with PBS, incubated with fluorescent secondary antibodies for 2 h in the dark at room temperature, washed three times with PBS, and stained with 4’,6-diamidino-2-phenylindole. Finally, they were observed under a fluorescence microscope.

### 2.13 Statistical Analysis

Data were statistically analyzed and plotted using GraphPad Prism 8 software. Data are expressed as mean ± standard deviation. One-way analysis of variance was used to analyze the differences between groups. A P-value <0.05 indicated statistical significance.

## 3. Results

### 3.1 In-Vivo Experiment Results

#### 3.1.1 Periodontal indexes

Compared with the C group, the GBI and PD increased in the P group (P<0.0001), confirming the successful construction of the periodontal disease model. Compared with the P group, the GBI (P<0.01) and PD (P<0.0001) decreased in the H group (Figure 1: D, E) (Table 3).

**Fig 1.**
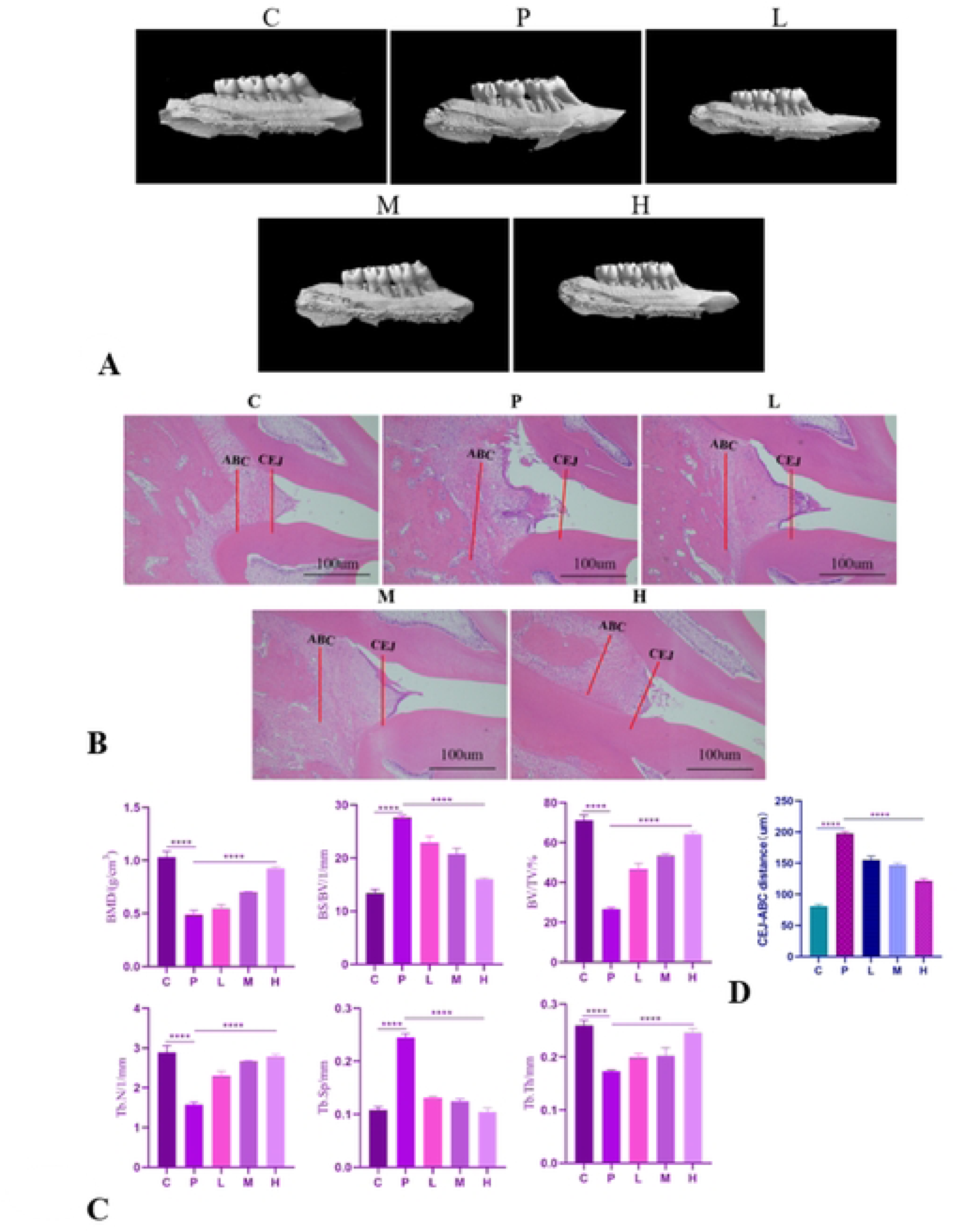
Periodontal index and alveolar bone structure. *A: Micro-CT reconstruction images of maxillary alveolar bone in rats of each group (n=4). *B: Observation of HE staining results in periodontal tissue of rats in each group. (CEJ is the enamel cementum and ABC is the crest of alveolar ridge, and the distance between them is the alveolar bone absorption.) (n=3). *C: Bone microstructure in the first molar region of maxilla. *D: CEJ-ABC distance (*p<0.05,**p<0.01, ***p< 0.001, ****p<0.0001)

**Table 2.**
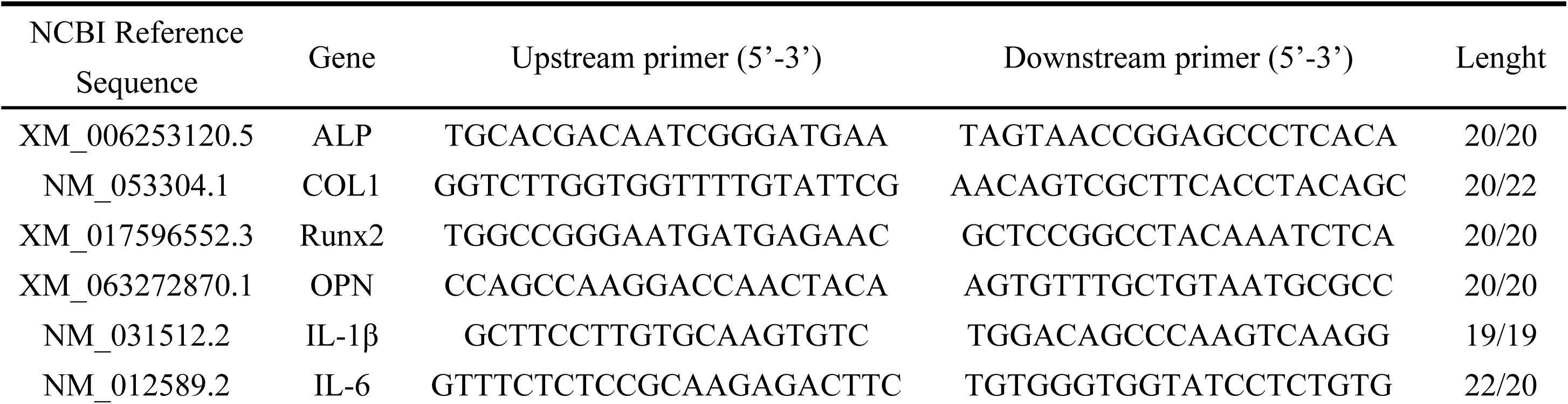
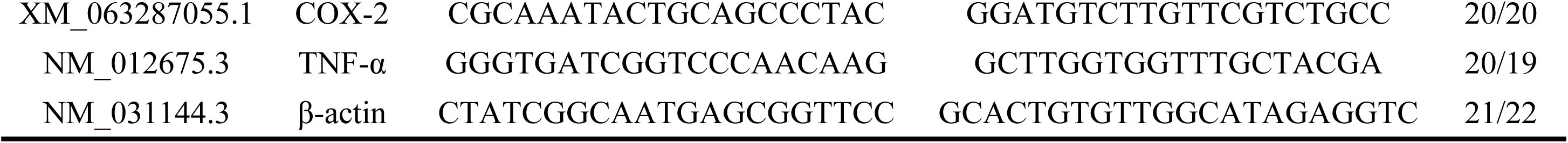
qRT-PCR primer sequences.

**Table 3.** The results of the GBI and PD.

| Periodontal index | GBI | PD (mm) |
| --- | --- | --- |
| C | 0.60±0.52<sup>b</sup> | 0.21±0.05<sup>c</sup> |
| P | 2.80±0.42 | 0.90±0.17 |
| L | 2.22±0.44 | 0.76±0.13 |
| M | 1.91±0.54 | 0.69±0.15 |
| H | 1.20±0.63<sup>a</sup> | 0.41±0.15<sup>b</sup> |

#### 3.1.2 Micro-CT Results

Compared with the C group, the P group demonstrated increased alveolar bone resorption, particularly in the exposure area of the first and second molar roots. The bone mineral density, bone volume fraction, trabecular number, and trabecular thickness in the region of interest of the maxillary alveolar bone were lower in the P group than in the C group (p<0.0001). Contrarily, the bone surface-to-bone volume ratio and trabecular separation were higher in the C group, confirming the establishment of the periodontal disease model. The H group bone mineral density, bone volume fraction, trabecular number, and trabecular thickness in the region of interest of the maxillary alveolar bone demonstrated significant improvements in various indicators, compared with the P group (P<0.0001) (Fig 1: A, C).

#### 3.1.3 H&E Staining Results

Fig 1 (B) illustrates histological examination results of periodontal tissues stained with H&E. The C group demonstrated intact alveolar bone without any signs of inflammation or damage to the gingival tissue. In contrast, the P group demonstrated the loss of gingival papillae, infiltration of inflammatory cells, and alveolar bone resorption, along with decreased alveolar crest height. The alveolar ridge height in the H group was significantly increased than that in the P group (Fig 1: D).

#### 3.1.4 IHC Staining Results

Compared with the C group, the P group demonstrated reduced levels of COL1 and Runx2 proteins (p<0.0001). The H group demonstrated higher protein expressions than the P group (p<0.0001) (Fig 2: A, B, and E) (Table 4).

**Fig 2.**
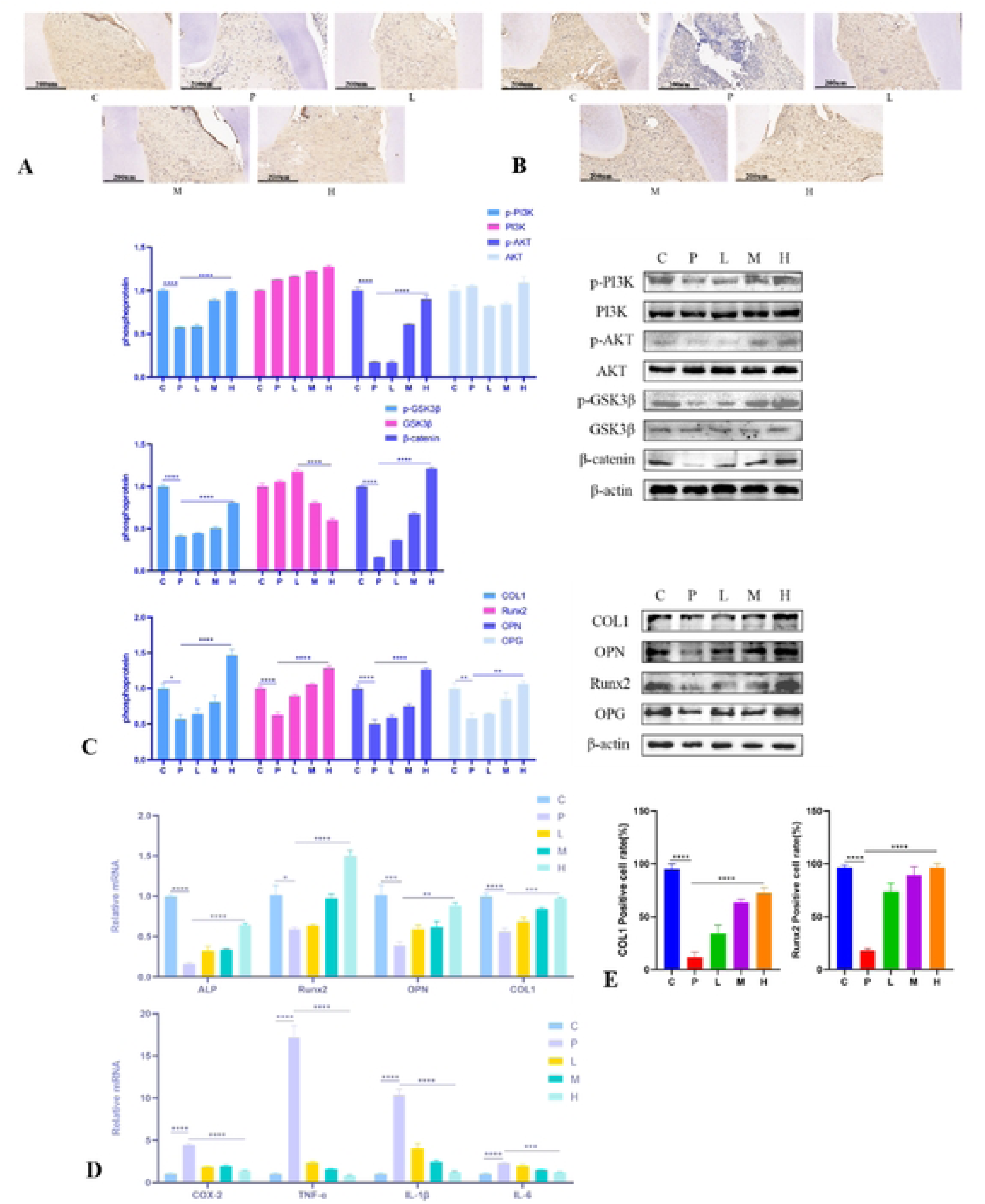
The expression results of proteins and mRNA in rat gingival tissue. *A: Runx2 expression level (n=3). *B: COL1 expression level (n=3). *C: the expression of proteins in gingival tissue of rats in each group (n=3). *D: Relative expression of osteogenic protein and inflammatory factors mRNA in gingival tissue of rats in each group (n=3). *E: Quantitative analysis of COL1 and Runx2 expression levels. (*p<0.05,**p<0.01, ***p<0.001, ****p<0.0001).

**Table 4.** quantitative IHC analysis results.

| Positive cell rate | COL1 (%) | Runx2 (%) |
| --- | --- | --- |
| C | 95.30±4.45<sup>d</sup> | 96.35±2.35<sup>d</sup> |
| P | 11.81±4.48 | 17.98±2.03 |
| L | 34.14±7.61 | 73.61±8.21 |
| M | 63.76±2.64 | 89.32±7.70 |
| H | 72.17±5.27<sup>d</sup> | 96.34±3.67<sup>d</sup> |

a: $p < 0.05$ compared to the P group; b: indicates $p < 0.01$ compared to the P group; c: indicates $p < 0.001$ compared to the P group; d: indicates $p < 0.0001$ compared to the P group.

#### 3.1.5 Western Blot Results

Compared with the C group, the P group demonstrated decreased expression of p-PI3K (p<0.0001), p-AKT (p<0.0001), p-GSK-3β (p<0.0001), β-catenin (p<0.0001), and osteogenesis-related proteins, such as OPN (p<0.0001), OPG (p<0.01), COL1 (p<0.05), and Runx2 (p<0.0001). Compared with the P group, the H group demonstrated increased expression of p-PI3K (p<0.0001), p-AKT (p<0.0001), p-GSK-3β (p<0.0001), β-catenin (p<0.0001), and osteogenesis-related proteins, such as OPN (p<0.0001), OPG (p<0.01), COL1 (p<0.0001), and Runx2 (p<0.0001) (Fig 2: C).

#### 3.1.6 qRT-PCR Results

Compared with the C group, the P group exhibited reduced expression of osteogenesis-related genes, including COL1A1 (p<0.0001), OPN (p<0.001), Runx2 (p<0.05), and ALP (p<0.0001). The H group exhibited increased gene expression than the P group, including COL1A1 (p<0.001), OPN (p<0.01), Runx2 (p<0.0001), and ALP (p<0.0001) (Fig 2: D).

### 3.2 Cell Experiment Results

#### 3.2.1 Effects of Carvacrol on Osteoblast Viability

Carvacrol cytotoxicity was assessed using the CCK-8 assay. Carvacrol concentrations of 50, 100, and 150 μM did not exert toxic effects on the cells. By contrast, concentrations ≥200 μM decreased the cell viability below 90%. Therefore, 50, 100, and 150 μM of carvacrol were used for subsequent experiments (Fig 3: E).

**Fig 3.**
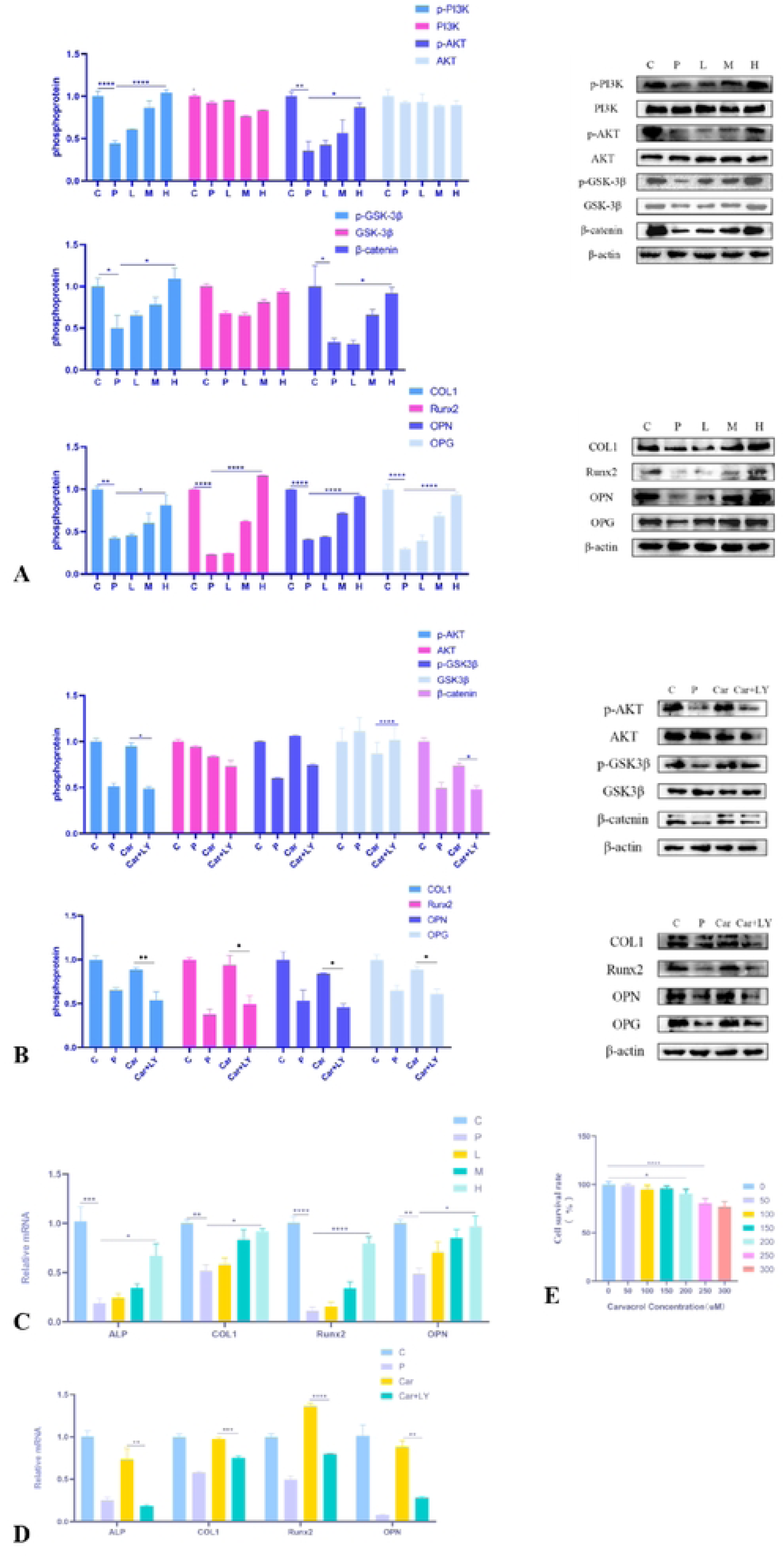
The expression results of proteins and mRNA in osteoblasts. *A: Expression of pathway-related proteins in cells of each group (n=3). *B: Expression of pathway-related proteins in cells of each group after adding LY294002 (n=3). *C: Relative expression of osteogenic protein mRNA in osteoblast (n=3). *D: Effects of LY294002 on Osteogenic Protein mRNA expression (n=3). *E: The proliferation of cells in different groups was observed by CCK-8. (*p<0.05,**p<0.01, ***p<0.001, ****p<0.0001).

#### 3.2.2 Western Blot Results

Compared with the C group, the P group demonstrated decreased expression of pathway proteins, such as p-PI3K (p<0.0001), p-AKT (p<0.01), p-GSK-3β (p<0.05), β-catenin (p<0.05), and osteogenesis-related proteins, including OPN (p<0.0001), OPG (p<0.0001), COL1 (p<0.01), and Runx2 (p<0.0001). Compared with the P group, the H group demonstrated increased expression of p-PI3K (p<0.0001), p-AKT (p<0.0001), p-GSK-3β (p<0.0001), β-catenin (p<0.0001), and osteogenesis-related proteins, such as OPN (p<0.0001), OPG (p<0.01), COL1 (p<0.0001), and Runx2 (p<0.0001) (Fig 3: A).

The Car+LY group demonstrated decreased expressions of p-AKT (p<0.05), p-GSK-3β (p<0.0001), β-catenin (p<0.05), and osteogenesis-related proteins, such as OPN (p<0.05), OPG (p<0.05), COL1 (p<0.001), and Runx2 (p<0.05), compared with the Car group (Fig 3: B).

#### 3.2.3 qRT-PCR Results

Compared with the C group, the P group exhibited reduced expressions of COL1A1 (p<0.01), OPN (p<0.01), Runx2 (p<0.0001), and ALP (p<0.001) genes. The H group exhibited increased gene expression than the P group, including COL1A1 (p<0.05), OPN (p<0.05), Runx2 (p<0.0001), and ALP (p<0.05) (Fig 3: C). The Car+LY group exhibited decreased expressions of ALP (p<0.01), OPN (p<0.01), COL1 (p<0.001), and Runx2 (p<0.0001) genes, compared with the Car group (Fig 3: D).

#### 3.2.4 Effects of Carvacrol on Osteoblast Differentiation

High-dose (150 μM) carvacrol increased ALP content and promoted the formation of mineralized nodules. However, LY294002 inhibited this positive effect (Fig 4: A, B, C, and D).

**Fig 4.**
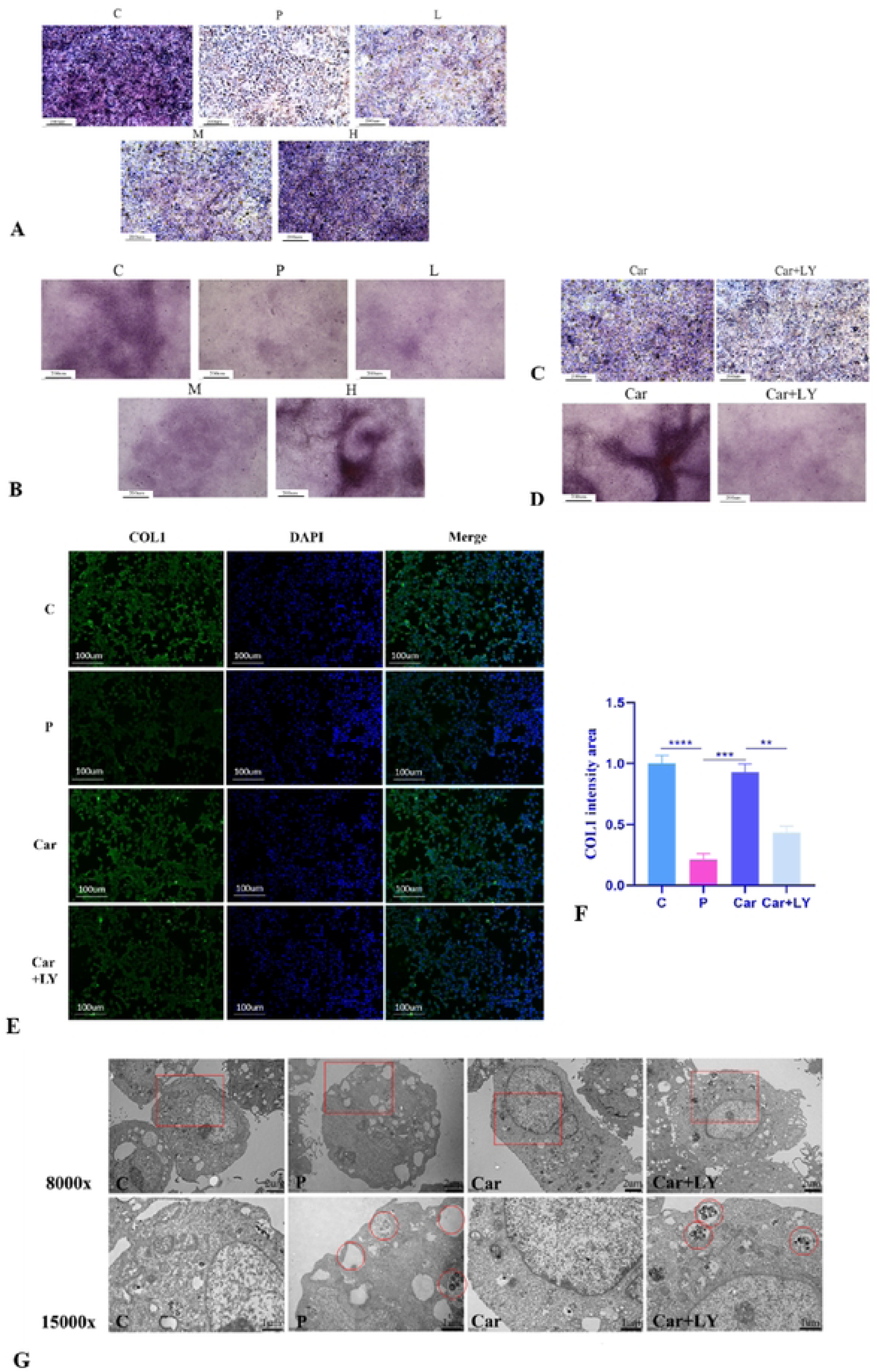
ALP and Alizarin Red staining, as well as IF and TEM result. *A and C: ALP staining after 7d (n=3). *B and D: Alizarin Red S Staining after 28d (n=3). *E: COL1 positive is green fluorescence (n=3). *F: Quantitative analysis of COL1 expression level (n=3). *G: Apoptotic bodies (red circles) were found in the cells (n=3) (15000x). (*p<0.05,**p <0.01, ***p<0.001, ****p<0.0001).

#### 3.2.5 IF Results

IF detection suggested that the green fluorescence intensity was comparable in the Car and C groups and higher than that in the P and Car+LY groups. Thus, carvacrol enhanced COL1 expression in osteoblasts, whereas LY294002 inhibited this promoting effect (Fig 4: E, F).

#### 3.2.6 TEM Results

TEM observations suggested that the endoplasmic reticulum swelled and apoptotic bodies formed during apoptosis. The P and Car+LY groups demonstrated a noticeably swollen endoplasmic reticulum with apoptotic bodies, whereas the Car group demonstrated improved symptoms (Fig 4: G).

## 4. Discussion

Periodontal disease is one of the most prevalent inflammatory conditions, characterized by bone and attachment destruction^[16]^. It can lead to gingival inflammation, gingival recession, bone mobility, and bone/tooth loss^[17]^. Conventional treatments primarily rely on antibiotics, such as metronidazole, which are central to inhibiting the growth of periodontal pathogens and controlling inflammation^[18]^. However, advances in medical research and evolving treatment philosophies suggest that merely suppressing bacterial growth and controlling inflammation are insufficient for comprehensively restoring periodontal tissue health. Additionally, overreliance on antibiotic treatments can lead to drug resistance^[19]^. The most ideal approach to treating periodontal disease is to inhibit the pathogenic bacteria responsible for the disease while repairing the alveolar bone. Under this treatment requirement, carvacrol, which possesses antibacterial, antioxidant, and osteogenic properties^[20]^, holds promise as a novel therapeutic agent for periodontal disease.

Carvacrol, a natural component of essential oils, has been widely studied because of its antibacterial, antioxidant, anti-inflammatory, anti-apoptotic, and bone-protective properties^[21]^. Carvacrol exhibits excellent antibacterial activity against periodontal pathogens^[6]^. Additionally, it is widely available, inexpensive, and holds great promise for further research. Building on this prior research, the current study demonstrated the role of carvacrol in regulating osteoblasts in alveolar bone affected by periodontal disease and explored its potential applications in periodontal therapy.

In this study, periodontal disease was successfully induced in the P group using ligature and LPS, leading to alveolar bone loss. The local application of varying concentrations of carvacrol hydrogel for 4 weeks improved periodontal disease symptoms, with the H group demonstrating substantial alveolar bone repair than the P group. Thus, the local application of carvacrol hydrogel effectively treats periodontal disease in rats and can repair the alveolar bone. This study confirms the role of carvacrol in promoting osteogenic protein expression. Thus, based on previous results, carvacrol may exert bone-repairing and regenerative effects through dual mechanisms—by inhibiting osteoclast differentiation and promoting osteoblast differentiation.

The PI3K/AKT signaling pathway promotes cell survival and reduces apoptosis. This pathway is central to promoting bone remodeling and development. The PI3K/AKT signaling pathway is central to processes, such as cell proliferation, differentiation, and apoptosis. This pathway has been strongly associated with osteogenic differentiation^[22]^. Therefore, if carvacrol can regulate this pathway, it may promote the osteogenic differentiation of osteoblasts. Xie et al.^[23]^ confirmed that carvacrol can enhance AKT phosphorylation. In this study, the phosphorylation level of the PI3K/AKT pathway decreased after LPS intervention; however, it increased after carvacrol treatment. The wingless-type MMTV integration site family canonical signaling pathway is associated with osteogenesis, β-catenin is a key protein in this pathway and can be degraded by GSK-3β^[24]^. Phosphorylation of the PI3K/AKT pathway phosphorylates GSK-3β, preventing β-catenin degradation and facilitating its accumulation in the nucleus, thereby promoting bone repair and regeneration. OPG, an osteoprotective factor, inhibits the binding of receptors involved in osteoclast pathways, thereby reducing osteoblast apoptosis while promoting their proliferation^[25]^. After blocking the PI3K/AKT pathway with LY294002, the therapeutic effects of carvacrol were partially inhibited. OPG expression decreased in the Car+LY group, compared with the Car group, weakening osteoclast inhibition and promoting osteoblast apoptosis. Furthermore, this inhibition reduced the expressions of COL1, OPN, and Runx2.

Western blot analysis demonstrated that the relative expressions of p-AKT, β-catenin, and osteogenesis-related proteins were lower in the Car+LY group than in the treatment group. Osteoblasts secrete calcium during the later stages of differentiation, forming calcified nodules^[26]^. The number of calcified nodules indicates osteogenic capacity. ALP—a marker of osteoblast activity—facilitates evaluating the osteogenic capacity^[27]^. ALP expression serves as an early indicator of bone formation, playing a major role in the early stages of mineralization in osteoblasts^[28]^. Therefore, ALP and Alizarin Red S staining kits were used to color the osteoblasts stimulated by different drug concentrations. Alizarin Red S staining can detect the calcified nodules secreted by mature osteoblasts, validating their osteogenic ability. In the Car+LY group, the intensity of ALP staining was lower than in the treatment group, and Alizarin Red S staining indicated fewer calcified nodules. When LY294002 was used to block the PI3K/AKT/GSK-3β pathway, it significantly inhibited the osteogenic effect of carvacrol. Therefore, the osteogenic effect of carvacrol is mediated through the regulation of the PI3K/AKT/GSK-3β signaling pathway. Alveolar bone repair is a complex biological process involving the proliferation and differentiation of osteoblasts. Because carvacrol promotes the osteogenic differentiation of osteoblasts, it may contribute positively to alveolar bone repair.

Both in vivo and in vitro results indicate that carvacrol hydrogel effectively reduced the inflammatory response associated with periodontal disease. The H group demonstrated the most profound alveolar bone repair effect. Carvacrol promotes the osteogenic differentiation of rat osteoblasts; the expression of the PI3K/AKT pathway in the H group closely matched that in the C group. After adding LY294002, the mRNA levels of inflammatory factors were higher in the Car+LY294002 group than in the treatment group. TEM results suggested that after LPS stimulation, the P group exhibited a marked inflammatory response, characterized by expanded endoplasmic reticulum and apoptotic bodies. After carvacrol treatment, apoptosis levels were lower in the H groups than in the P group. Thus, carvacrol effectively reduces osteoblast apoptosis. In this study, in vivo and in vitro co-verification methods were adopted; however, in vitro experiments cannot completely simulate the in vivo state, which may present some limitations.

## 5. Conclusion

In summary, carvacrol exerts therapeutic effects in periodontal disease by activating the PI3K/AKT/GSK-3β pathway and promoting osteoblast proliferation and differentiation. Moreover, it reduces osteoblast apoptosis and the expression of inflammatory factors. These findings provide a theoretical foundation for the application of carvacrol hydrogel in periodontal disease treatment.

## 6. Acknowledgments

We especially thank the Key Laboratory of the Provincial Education Department of Heilongjiang for Common Animal Disease Prevention and Treatment. This project is supported by the “Program for Young Talents of Basic Research in Universities of Heilongjiang Province (YQ2023C016)”.

